# Selectomic and Evolvability Analyses of the Highly Pathogenic Betacoronaviruses SARS-CoV-2, SARS-CoV, and MERS-CoV

**DOI:** 10.1101/2020.05.05.078956

**Authors:** Meghan May, Bahman Rostama, Ryan F. Relich

**Affiliations:** Department of Biomedical Sciences, College of Medicine, University of New England, Biddeford, ME USA; Department of Pathology and Laboratory Medicine, Indiana University School of Medicine, Indianapolis, IN USA

**Keywords:** COVID19, Coronavirus, Evolvability, Evolution, Emerging Viruses, Emerging Infectious Diseases, Diversifying Selection, SARS-CoV-2, SARS-CoV, MERS-CoV

## Abstract

SARS-CoV-2, the causative agent of COVID-19, is widespread in several countries around the world following its late-2019 emergence in the human population. Rapid development of molecular diagnostic tests and subunit vaccines have been prioritized, and as such evaluating the SARS-CoV-2 genomic plasticity and evolutionary dynamics is an urgent need. We determined the SARS-CoV-2 selectome by calculating rates of pervasive and episodic diversifying selection for every amino acid coding position in the SARS-CoV-2 genome. To provide context for evolutionary dynamics of a highly pathogenic betacoronavirus following a zoonotic spillover into human hosts, we also determined the selectomes of SARS-CoV and MERS-CoV, and performed evolvability calculations for SARS-CoV-2 based on SARS-CoV. These analyses identify the amino acid sites within each coding sequence that have been subjected to pervasive diversifying selection or episodic diversifying selection, and report significantly evolvable sites in the ORF1a polyprotein, the spike protein, and the membrane protein of SARS-CoV-2. These findings provide a comprehensive view of zoonotic, highly pathogenic betacoronavirus evolutionary dynamics that can be directly applied to diagnostic assay and vaccine design for SARS-CoV-2.

## Introduction

As of December 2019, seven coronaviruses have been identified as causes of human respiratory tract infections. Four of these viruses, the human coronaviruses (HCoVs) HCoV-229, HCoV-OC43, HCoV-NL63, and HCoV-HKU1, co-circulate annually with many other respiratory viruses across the globe during colder months, and they are usually associated with mild upper and/or lower respiratory tract infections. Severe diseases can be caused by HCoVs, and fatal infections have been documented, especially among individuals with compromised immune systems, young children, and older adults (1–4). In late 2002, numerous cases of severe viral pneumonia were reported in Guangdong Province, China. Over the following nine months, 8,448 total cases and 774 deaths were reported in 26 countries across the globe. This disease, called severe acute respiratory syndrome (SARS) was determined to be caused by a novel coronavirus named SARS coronavirus (SARS-CoV). SARS is characterized by fever, chills, headache, muscle aches, and other generalized symptoms (5). Severe complications varied in frequency among described patient cohorts, but most fatal cases were associated with severe pneumonia. The case fatality rate of SARS-CoV infection is approximately 10%; however, considerable differences in case fatality rates were observed in patients of different age groups (6). In 2012, a patient from Saudi Arabia with severe lower respiratory tract disease was diagnosed with a novel coronavirus, later called Middle East respiratory syndrome coronavirus (MERS-CoV) (7). As of November 2019, nearly 2,500 cases of Middle East respiratory syndrome (MERS) have been diagnosed in 12 countries, and the case fatality rate is estimated to be approximately 34% (8). MERS-CoV continues to cause cases, but they are infrequent and appear to be associated with dromedary camel-to-human spillover events followed by occasional human-to-human transmission events during caretaking (9).

At the end of 2019, a novel coronavirus called severe acute respiratory syndrome-related coronavirus 2 (SARS-CoV-2) emerged in Wuhan, China. The first cases were identified among individuals with links to a seafood market in Wuhan (10). The detection of similar viruses in wild-caught rhinolophid bats and pangolins in China suggests a possible zoonotic reservoir and/or intermediate host(s), respectively; however, recent data suggests that this virus did not directly spill over into the human population from pangolins (11). The exact source of this virus is not currently known, and the description of a small number of clades raise the possibility of a complex series of events preceding the spillover event (12). As of this writing, over 2,000,000 human cases of SARS-CoV-2-associated disease, called <u>co</u>ronavirus <u>di</u>sease 20<u>19</u> (COVID-19) have been reported from a total of 210 countries scattered across all continents except Antarctica (13). The case fatality rate (CFR) of COVID-19 varies, but the overall CFR is approximately 2 - 3%, and the disease appears to be most severe in individuals of advanced age. Interestingly, children seem to be spared from severe complications; however, fatal cases among pediatric patients have been described (14–16).

Coronaviruses leading to acute respiratory distress syndrome (ARDS; MERS-CoV, SARS-CoV, SARS CoV-2) entered the human population following zoonotic spillover events. These viruses are also known to infect animals, including bats (MERS-CoV, SARS-CoV, and SARS-CoV-2), dromedary camels (MERS-CoV), masked palm civets (SARS-CoV), non-human primates (MERS-CoV, SARS-CoV), ferrets (SARS-CoV-2), house cats (SARS-CoV-2), and rodents (MERS-CoV, SARS-CoV) (1, 17–18). Closely related coronaviruses have also been detected in animals, but they have yet to be implicated in human infections (11).

Coronaviruses possess the largest genomes of all RNA viruses so far described. They are comprised of monopartite, positive-sense, single-stranded RNA (+ssRNA) molecules that vary in length from 27-32 kbp (19). Genomes encode four structural proteins (spike, S; envelope, E; membrane, M, and nucleoprotein, N) and 16 nonstructural proteins (NSPs) encoded by a large open reading frame (ORF) termed ORF1ab (20). Initial attachment to host cells occurs following binding of the S protein to its cellular receptor, identified so far for the ARDS-associated viruses to include angiotensin-converting enzyme 2 (ACE2; SARS-CoV, SARS-CoV) and dipeptidyl peptidase 4 (DPP4; MERS-CoV) (20). Coronaviruses feature exonuclease activity that maintains fidelity during genome replication, which is an uncommon feature among RNA viruses (21). This unusually high genomic stability makes areas of plasticity in coronaviruses particularly noteworthy, and underscores the need to evaluate elevated rates of diversity carefully. Here we present a comprehensive selectomic and evolvability analysis of SARS-CoV-2, SARS-CoV, and MERS-CoV, the coronaviruses causing ARDS in humans, with particular emphasis on evaluating diversity and evolvability in diagnostic targets and sites associated with host adaptation.

## Highly Pathogenic Betacoronaviruses are Subject to both Pervasive and Episodic Diversifying Selection

Pervasive Diversifying Selection (PDS) was detected in the N, ORF1a, and ORF1b protein products for all three viruses (Figure 1A-1E, Table 1). The S and M proteins of SARS-CoV and MERS CoV also had multiple sites under PDS, including sites within the S protein binding domain (Fig 2F). A majority of sites under PDS that were able to be mapped onto three-dimensional protein structures were located in disordered regions [3/3 SARS-CoV-2; 7/7 SARS-CoV; 9/11 MERS CoV; Fig. 2A – 2G]. Predicted N-linked glycosylation sites did not appear to correlate with the position of sites under PDS (Fig. 1A-1E). Episodic Diversifying Selection (EDS) was detected in the ORF1a protein product for all three viruses. The S protein of SARS-CoV-2 and the N protein and ORF1b protein product of MERS-CoV also had several sites under EDS (Figure 1A-1E, Table 1). The majority of sites under EDS that were able to be mapped onto the three-dimensional protein structure of the SARS-CoV S protein were located in disordered regions [12/14; Fig. 2F]. In contrast, the only sites under EDS that were able to be mapped onto three-dimensional protein structures for MERS-CoV were located exclusively within β-sheet formations [3/3; ORF1a deubiquitinase domain; Fig. 2A]. The single site under EDS for SARS-CoV was not able to be mapped onto a predicted structure. Predicted N-linked glycosylation sites did not appear to correlate with the position of sites under EDS (Fig 1A-1E).

**Table 1:**
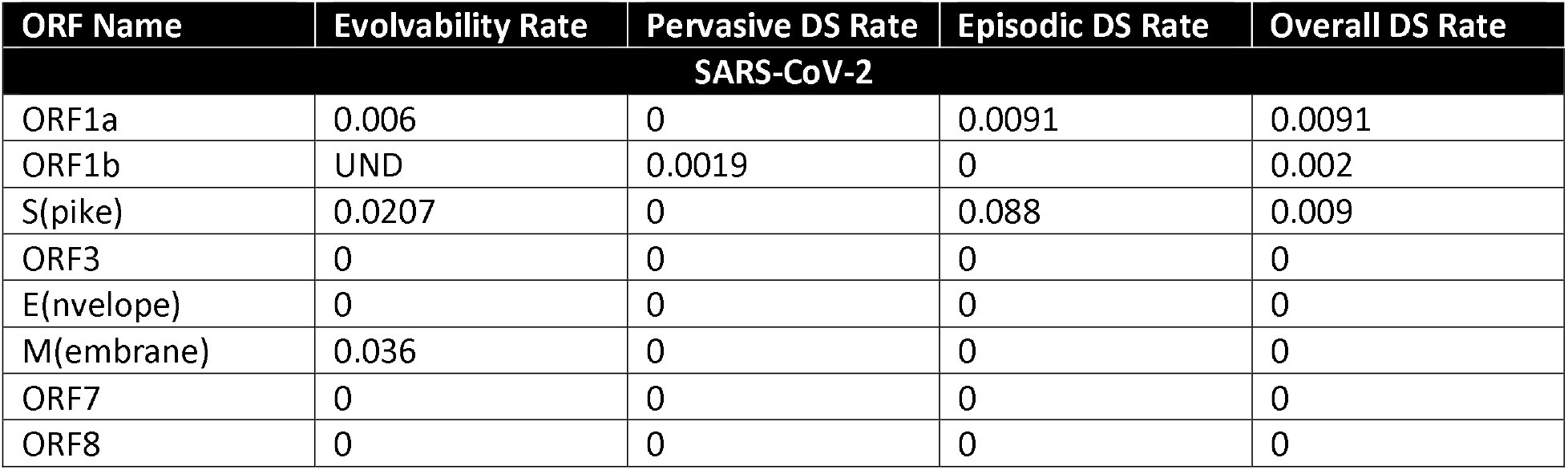
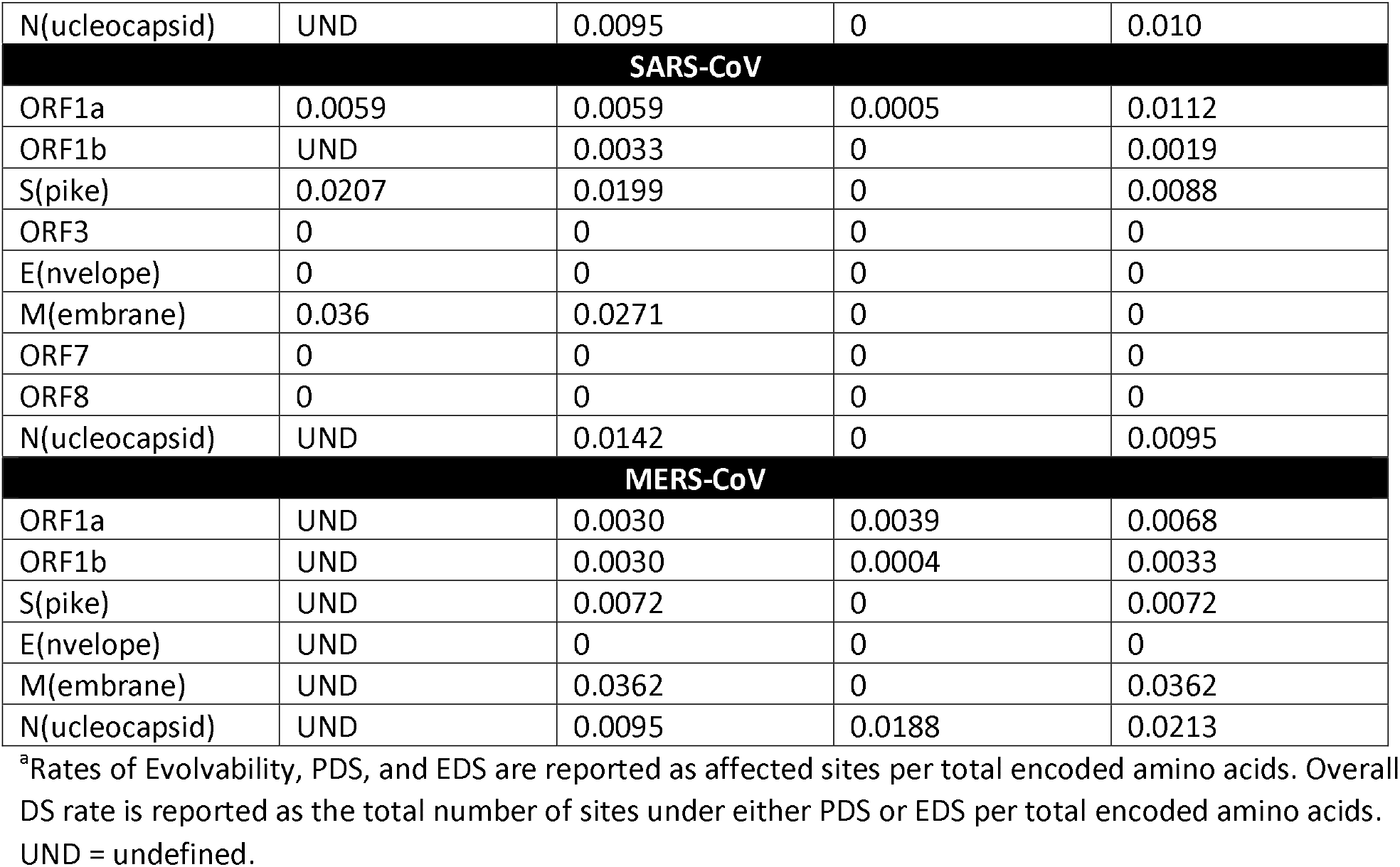
Diversity across Sites by Open Reading Frame^a^.

**Figure 1:**
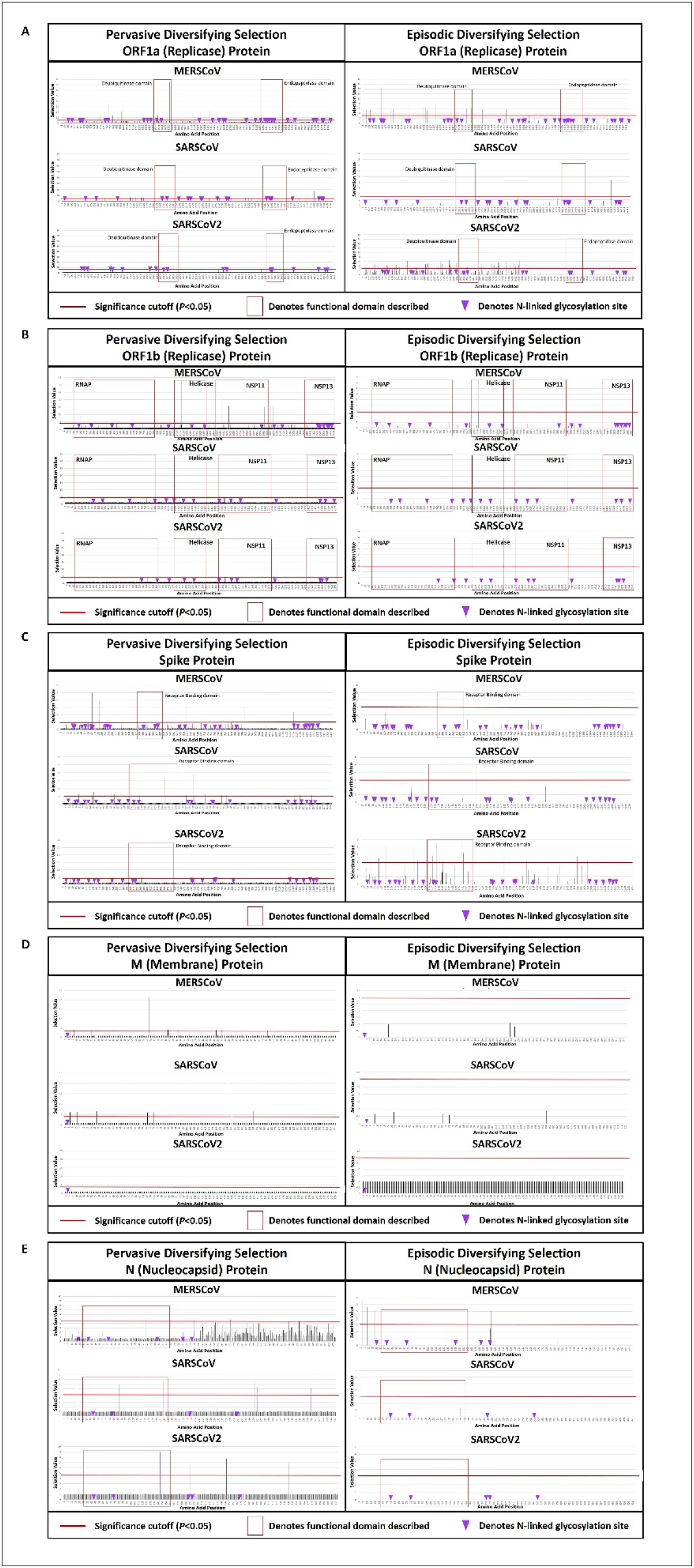
PDS and EDS Predictions in SARS-CoV-2, SARS-CoV, and MERS-CoV. Selection values (Y axis) for ORF1a (A), ORF1b (B), S (C), M (D), and N (E) are plotted at each amino acid position (X axis). PDS values for each trait are shown on the left side of the panel, and EDS values are shown on the right. The cutoff for significance is shown as a red line; values rising above the line indicate diversifying selection acting at that amino acid position. Important functional domains are boxed in red and identified on each panel. Predicted N-linked glycosylation sites are shown as purple arrowheads.

**Figure 2:**
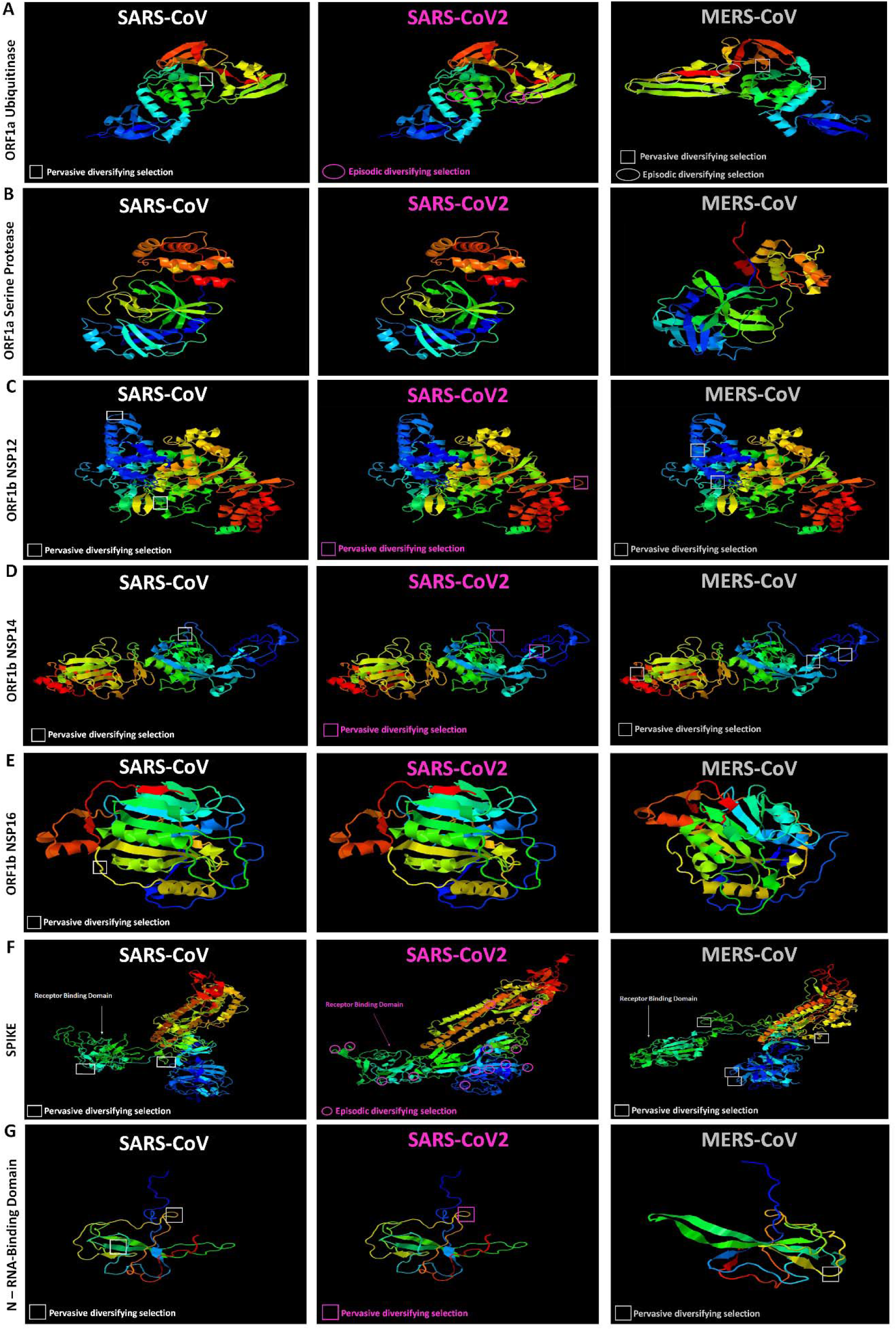
Structural Predictions of Positively Selected SARS-CoV-2, SARS-CoV, and MERS-CoV Proteins. Three dimensional models were generated for portions of the ORF1a polyprotein (A, deubiquitinase/papain-like protease; B, endopeptidase/serine protease), the ORF1b polyprotein (C, NSP12; D, NSP14; E, NSP16), S (F), and N (G, RNA-binding domain) of SARS-CoV-2, SARS-CoV, and MERS-CoV. Sites under PDS are boxed, and sites under EDS are circled. Selected sites shown in Figure 1 that are not located within the predicted regions are not shown.

PDS was detected in all three viruses, and was most frequently found in MERS-CoV. SARS-CoV had an intermediate level of PDS and very little EDS, whereas SARS-CoV-2 had three traits under PDS and two under EDS. As PDS reflects viral response to persistent selective pressure, the comparatively longer tenure of MERS-CoV as a human pathogen likely accounts for its higher frequency. Similarly, the circulation of SARS-CoV for at least 9 months gave it far longer to adapt to the human host environment via diversification than SARS-CoV-2 has yet experienced. In contrast, EDS is often seen transiently after a sudden shift in habitat, and tends to become less detectable over time as sites either optimize within the new host environment or become subject to PDS. It is notable that while the recently emerged SARS-CoV-2 has only two traits under EDS, the S protein EDS rate is the highest selection rate observed for any trait across the three viruses (Table 1). Elevated EDS occurring soon after a host species jump is highly consistent with the role of the S protein in mediating host cell attachment and fusion. SARS-CoV, which appeared once in the human population, circulated for 9 months, and has never re-emerged, had very little detectable EDS. This, too, is consistent with a single introduction into a human host followed by circulation within that host until sites begin to stabilize or operate under PDS. Three MERS-CoV traits were subject to both PDS and EDS. Given that MERS-CoV can be transmitted from human-to-human and from dromedary camels to humans, and has been causing infections in human hosts since at least 2012, the combination of both types of diversifying selection likely reflects simultaneous attempts to adapt to multiple host habitats while passing between them. The E, ORF3, ORF7, and ORF8 protein products were uniformly under purifying selection and were highly conserved.

Three-dimensional structural predictions were made for ORF1a, ORF1b, S, and N from each virus (Fig 2A – 2G), and any sites under PDS or EDS were noted. Sites under PDS across all three species exhibited a striking tendency to occur in disordered, low-complexity regions whose structures tend to be highly flexible (Fig. 2). The majority of sites under EDS that were able to be mapped onto the three-dimensional protein structure of the SARS-CoV-2 S protein were located in disordered regions [12/14; Fig. 2F]. In contrast, the only sites under EDS that were able to be mapped onto three-dimensional protein structures for MERS-CoV were located exclusively within β-sheet formations [3/3; ORF1a deubiquitinase domain; Fig. 2A]. It is conceivable that EDS acting on sites within a defined structure allows for fine-tuning of camel-versus-human host target specificity while maintaining deubiquitinase enzymatic function as MERS-CoV passes between the two host habitats. Predicted N-linked glycosylation sites did not appear to correlate with the position of sites under PDS or EDS for any trait of any of the three viruses, although structural analysis to confirm modifications of predicted sites is a critical measure to truly establish the absence of a connection between these factors.

## ORF1a, S, and M of SARS-CoV-2 Have Several Evolvable Amino Acid Sites

Evolvability could be calculated for the ORF1a protein product, the S protein, and the M protein. Exclusive purifying selection across SARS-CoV and SARS-CoV-2 prevented evolvability calculations for E, ORF3, ORF7, and ORF8, and pervasive diversifying selection in SARS-CoV-2 prevented them for ORF1b and N. Several significantly evolvable amino acid sites were identified in the ORF1a polyprotein, but the distribution was non-uniform. Proteins NSP1, the nucleic acid binding domain, the endopeptidase, and NSP9 had no evolvable sites, whereas NSP2, NSP3, NSP4, NSP7, NSP8, NSP10, the ADP ribose-binding domain, and the deubiquitinase (papain-like protease) domain had several evolvable sites (Fig 3A). Significantly evolvable sites were found throughout the S protein, including within the binding motif (Fig. 3B, Table 2). Of those that were able to be mapped, evolvable amino acid positions in S and ORF1a were frequently found in predicted disordered regions (Fig. 3A – 3B). Eight evolvable sites were identified in the M protein irrespective of predicted secondary structure, occurring within predicted α-helices, β-strands, and coiled coils (Fig. 3C).

**Table 2:** Evolvability and Diversity - Binding Domain and Subunit Vaccine Antigens^a^.

| CoV Species | Binding Domain | Binding Domain Diversity |  |  | Vaccine Antigen<sup>b</sup> Diversity |  |  |
| --- | --- | --- | --- | --- | --- | --- | --- |
|  |  | E | PDS | EDS | E | PDS | EDS |
| SARS-CoV-2 | 343 - 597 | 5 | 0 | 4 | 26 | 0 | 11 |
| SARSCoV | 317 - 564 | 5 | 6 | 0 | 26 | 25 | 0 |
| MERS-CoV | 375 - 500 | UND | 0 | 0 | UND | 9 | 0 |

<sup>a</sup>E = evolvable sites; UND = undefined
<sup>b</sup>Vaccine antigen includes the S protein sequence, as previously described for MERS-CoV and SARS-CoV (22 – 24)

**Figure 3:**
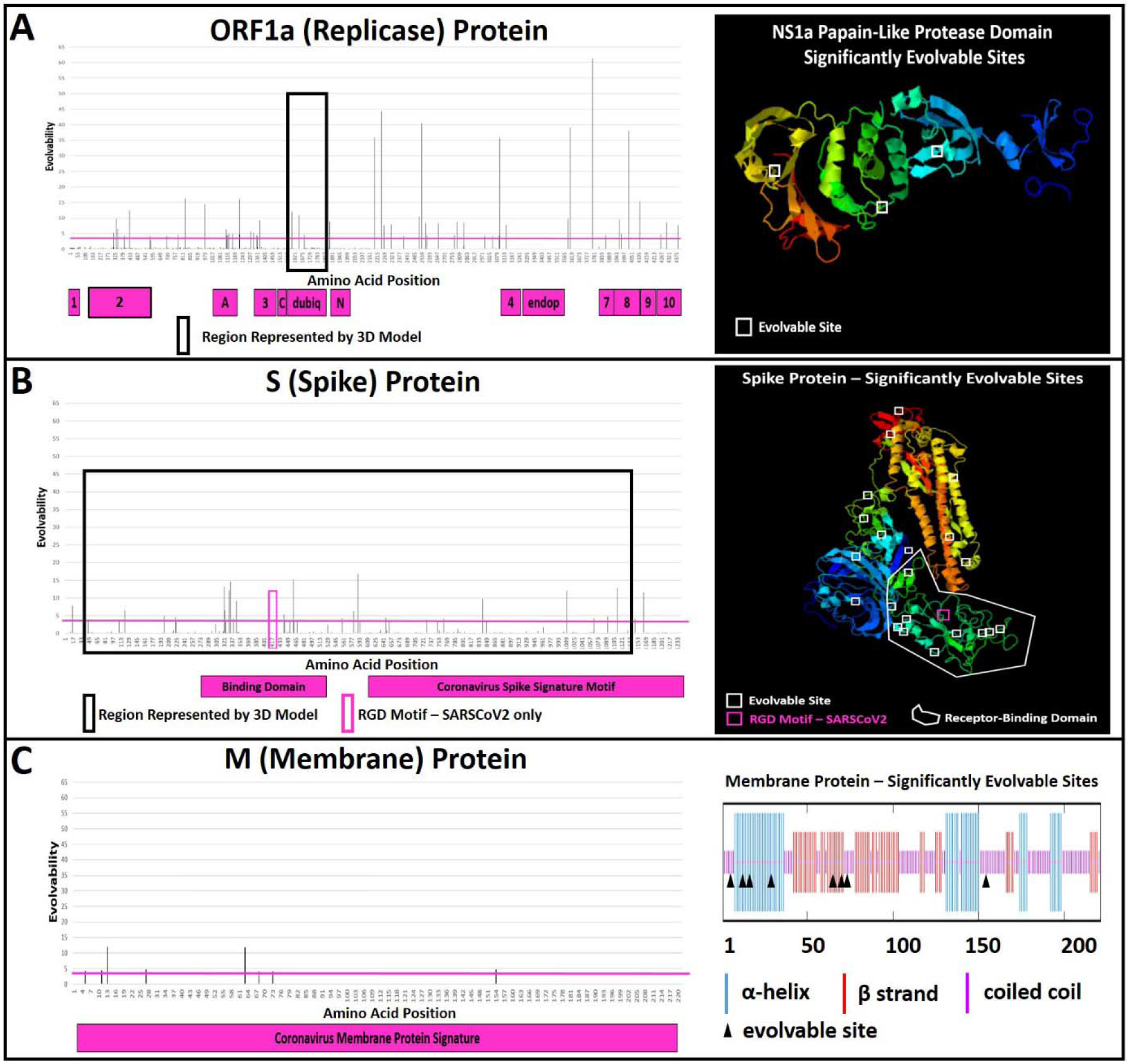
Evolvability Measures of SARS-CoVs. Evolvability (Y axis) was detected at multiple amino acid sites (X axis) in the ORF1a polyprotein (A), the S protein (B), and the M protein (C). The cutoff for significance is shown as a pink line; values rising above the line indicate that the amino acid position is significant evolvable. Black boxes indicate the region depicted in the accompanying three dimensional structural models (A, B). All amino acid positions are included in the M protein secondary structure prediction (C). Evolvable residues appearing within the structural predictions are shown as white boxes (A, B) or black arrowheads (C), and were notably frequent in disordered regions. The novel S protein R-G-D motif arising in SARS-CoV-2 is indicated as a pink box and appears within a disordered region of the binding domain (B). Functional domains are indicated as pink blocks below the X axis [label abbreviations - 1: NSP1; 2: NSP2; A: ADP-ribose binding domain; C: cleavage target; 3: NSP3; Dubiq: deubiquitinase (papain-like protease); N: nucleic acid binding domain; 4: NSP4; Endop: endopeptidase (serine protease); 7: NSP7; 8: NSP8; 9: NSP9; 10: NSP10].

Using the recently emerged, relatively low-diversity SARS-CoV-2 as a base and the extensively circulated, pervasively selected SARS-CoV as an endpoint query, the evolvability of each amino acid position for ORF1a, S, and M was calculated. Uniform purifying selection acting on ORF3, E, ORF7, and ORF8 in SARS-CoV and SARS-CoV-2 indicate that these traits are not significantly evolvable following a host species jump. In contrast, PDS has already measurably acted on ORF1b and N, making SARS-CoV-2 an already inappropriate base point for these traits. Numerous sites in the ORF1a protein product and the S protein are significantly evolvable, including some within known functional motifs. Evolvable sites were detected in NSP2, NSP3, NSP4, NSP7, NSP8, NSP10, the ADP ribose-binding domain, and the deubiquitinase (papain-like protease) domain, suggesting that the functions of these ORF1a cleavage-product proteins are subject to diversification during adaptation to a human host. Similarly, many evolvable sites were detected within the S protein including within the binding domain. The majority of these sites also fall within highly disordered regions (Fig. 3B). This is consistent with the notion of host cell binding and entry being a strong driver of diversification upon entry into a new host. Further, the role of the S protein as an immunodominant antigen for both SARS-CoV (28) and MERS-CoV (29) suggests that, in addition to functional selection, immune selection could also play a role in diversification and evolvability. Notably, a SNP found across all SARS-CoV-2 strains reported results in a change from lysine in SARS-CoV to arginine in SARS-CoV-2. This substitution is moderately synonymous (K-R BLOSUM62 score = 2), and therefore the site is not under significant PDS. While this is also not flagged as a significantly evolvable site, it nevertheless introduces an R-G-D motif in the S protein of SARS-CoV-2, which may confer integrin-binding capacity onto the virus (30). The biological impact of this change has yet to be explored.

## SARS-CoV-2 Diagnostic Tests and Subunit Vaccines Target Positively Selected, Evolvable Traits

Management and control of the COVID-19 pandemic requires rapid and expansive diagnostic testing. The diagnosis of coronavirus infections including COVID-19 is currently performed by nucleic acid amplification test (NAAT) methods such as real-time reverse transcription PCR (rRT-PCR). Several NAATs to detect SARS-CoV-2 have received Emergency Use Authorization from the United States Food and Drug Administration and equivalent international authorities (Table 3). The majority of these assays include primers and probes targeting multiple sites within the SARS-CoV-2 genome; however, it is notable that no tests exclusively target ORFs with evolvability and/or diversifying selection rates of 0. Only four SARS-CoV-2 NAAT platforms (*i.e.*, the BioFire 2.0, the Cepheid Xpert Xpress SARS-CoV-2, the Roche cobas SARS-CoV-2, and the Luminex NxTAG CoV Extended Panel) include a target ORFs (ORF8, E, E, and E, respectively) that is under purifying selection and does not have significantly evolvable sites. While the specific primer and probe sequences of the assays described in Table 3 are largely proprietary and thus could not be assessed in this study, the locations of those sequences and of future diagnostic tests should be evaluated in the context of sites encoding significantly evolvable amino acid positions and those under pervasive diversifying selection to minimize the chance of divergent viruses escaping detection (Supplemental Datasets S1, S2, S3, S4). Similarly, proposed nucleic acid and subunit vaccines must similarly consider evolvable and positively selected sites to minimize the possibility of viral escape variants.

**Table 3:**
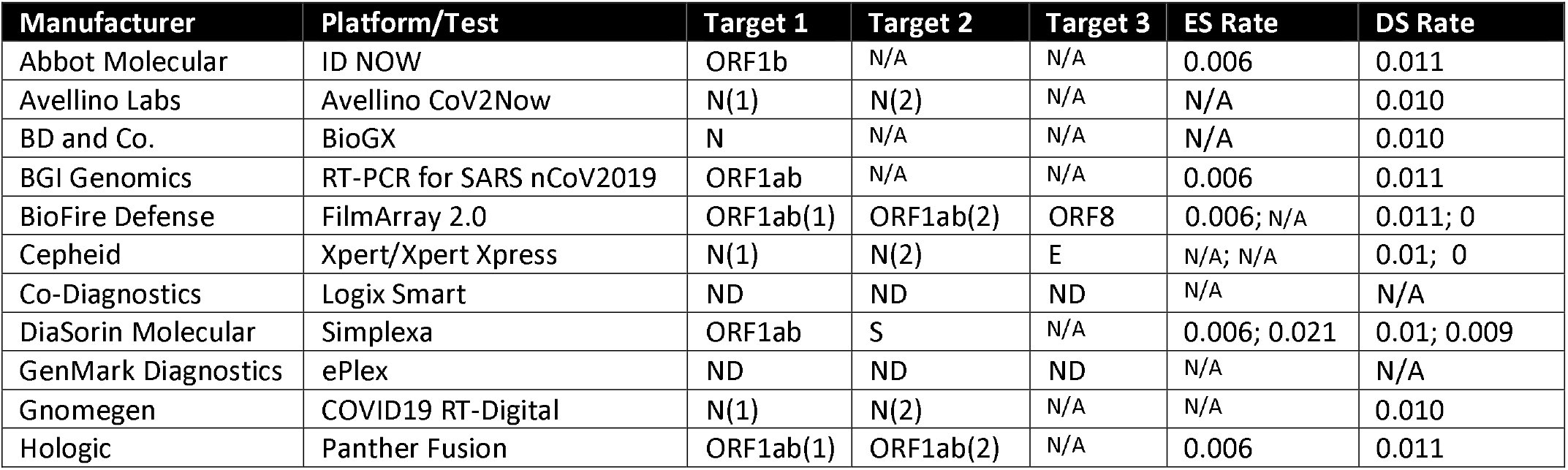
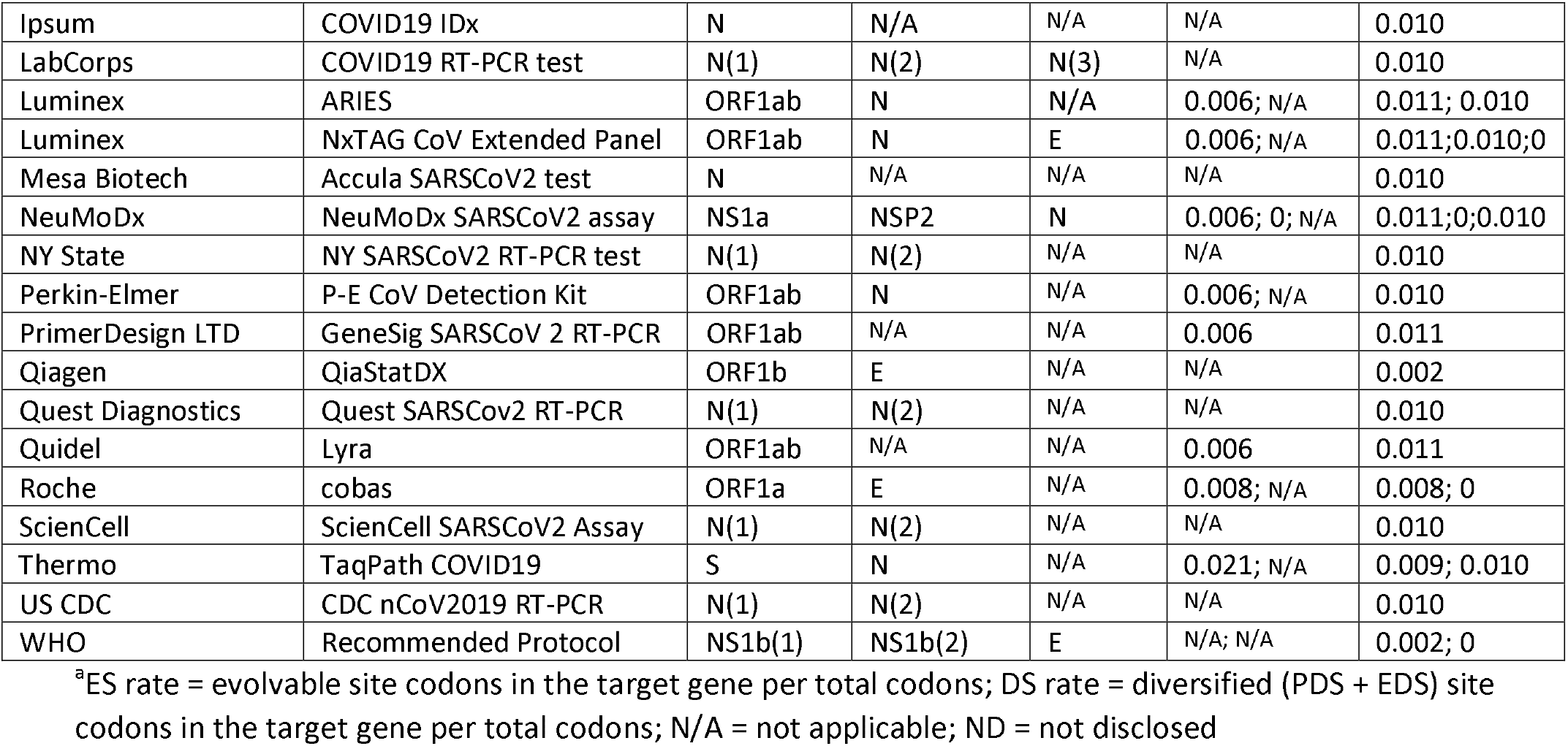
SARS CoV-2 Diagnostic Test Targets^a^.

## Discussion

The novel coronavirus SARS-CoV-2 is the third *Betacoronavirus* to emerge in the past twenty years. All known human-pathogenic coronaviruses are transmissible from human to human and are also known to infect animals (1, 17, 25). Other coronaviruses, including SARS-CoV-like coronaviruses (SARS-CoVs), have also been detected in animals, but they have yet to be implicated in human infections (11). Many comparative genomic analyses have described single nucleotide polymorphisms (SNPs) among clinical isolates of SARS-CoV-2 (12, 26), but ascribing function to those SNPs absent of context can lead to over-interpretation (27). Determination of the sites under diversifying selection and sites that are significantly evolvable in those coronaviruses causing ARDS (*i.e.*, SARS-CoV, MERS-CoV) that have circulated among a novel human host can inform expectations around accumulating diversity in SARS-CoV-2 as it begins the process of adaptation to a human habitat.

Diversification of viral adhesins is a common mechanism of host adaption or expansion of target tissues, and PDS (MER-CoV, SARS-CoV) and EDS (SARS-CoV-2) acting on sites in the S protein is consistent with that trend. However, we detected several additional traits with sites that are diversifying in response to selective pressure, underscoring the need for multiple areas of the virus infectious cycle to adapt to a novel host for a successful species jump to occur. For example, sites under PDS were detected in the N protein of all three viruses, possibly indicating that interactions between nucleocapsids and host cell ER-Golgi intermediate compartment (ERGIC) as progeny virions mature is a point where host cell permissiveness can exert itself. Similarly, PDS in the nucleic acid binding domain and the M protein may reflect that RNA targets or components needed for envelope maturation represent points critical for novel host adaptation.

Identification of evolvable residues is possible by the presence of homologues in two species, one of which is under PDS and the other of which is not in its current habitat (31). Human coronaviruses can be both plastic (*e.g.*, HCoV-OC43 [32]) and highly conserved (*e.g.*, HCoV-229E [33]), indicating that evolvability analysis is a technique with potential to identify more mutable species in wildlife reservoirs. This underscores the need to conduct continuous viromic surveillance of wildlife reservoirs, as identification of coronaviruses with significantly evolvable traits are likely at higher probability for successful zoonotic spillover events.

Here we describe the positive selectomes of SARS-CoV-2 and SARS-CoV, the partial positive selectome of MERS-CoV, and the identification of significantly evolvable sites in SARS-CoV-2 based on the natural history of SARS-CoV. Pervasive diversifying selection was detected in the significantly evolvable ORF1a polyprotein and the S protein of MERS-CoV and SARS-CoV, and EDS was detected in the same proteins of SARS-CoV-2. These findings suggest that host adaptation of SARS-CoV-2 to humans is likely to lead to viral diversification parallel to that seen for SARS-CoV and MERS-CoV. Detection of PDS in all three species in the ORF1b polyprotein and N protein are further consistent with that outcome. Consideration of significantly evolvable residues identified can inform the design and refinement of diagnostic tests and subunit vaccines for SARS-CoV-2, and periodic revisions to the SARS-CoV-2 selectome over time will be immensely valuable to confirm the predicted evolvability of noted amino acid sites.

## Supporting information

MERS-CoV Strain Information

SARS-CoV Strain Information

SARS-CoV-2 Strain Information

Data Supplement 1 - SARS-CoV-2 selection values

Data Supplement 2 - SARS-CoV selection values

Data Supplement 3 - MERS-CoV selection values

Data Supplement 4 - Evolvability Values

## Author Contributions

MM and RFR conceived of the project and designed the study; BR assembled datasets and strain information documentation; MM and BR annotated genomes and performed computational analyses; RFR assembled diagnostic target sets; MM drafted the initial manuscript; MM, BR, and RFR revised and edited the final manuscript.

## Data Availability

All data relevant to this manuscript are presented in the main text and figures, are available in the Supplemental Materials, are posted at doi: 10.5281/zenodo.3756937, or can be accessed in public databases as noted in Supplemental Tables S1 – S3.

## Acknowledgements

We gratefully acknowledge all the authors who have shared their genome data by depositing on GISAID (35) or in GenBank (36, 37). A table with full author lists for all SARS-CoV-2 sequences used can be found here: doi: 10.5281/zenodo.3756937.

