## Supplemental Materials for "Selectomic and Evolvability Analyses of the Highly Pathogenic Betacoronaviruses SARS-CoV-2, SARS-CoV, and MERS-CoV"

1.) **Materials and Methods**

*Coronavirus Species and Strains*. Strains of SARS-CoV-2, SARS-CoV, and MERS-CoV with publicly accessible complete genomes were selected and accessed through GISAID (35) and GenBank (36, 37). A minimum of 3 distinct sequences is required to perform evolutionary selection analyses, and additional strains increase the resolution and statistical power of each analysis. To that end, our dataset consisted of 14 SARS-CoV strains (2003 – 2004, 9 months), 30 MERS-CoV strains (10/30/2012 – 10/20/2018, 6 years), and 55 SARS-CoV-2 strains (12/29/2019 – 01/29/2020, 4.5 weeks). Genomes accessed as .FASTA nucleotide sequences were annotated using NCBI ORFfinder (37) followed by BLAST (38). The biological source, geographic location, approximate date of isolation, database accession numbers, and depositing authors for each strain are noted in Supplemental Tables S1 – S3, and at doi: 10.5281/zenodo.3756937.

*Evolutionary Selection Analysis.* Two distinct methods for detecting evolutionary selection were employed. The first was a Bayesian, fixed effects model to detect pervasive positive (diversifying) and pervasive negative (purifying) selection (PDS and PPS, respectively) rates at each site, and assumes consistent selective pressures are operating on all strains. Calculations were performed using the Fast, Unconstrained Bayesian AppRoximation (FUBAR) model (39) on the DataMonkey 2.0 server (40). The Bayes factor calculated for each site represents the likelihood ratio of the marginal ratio for agreement between two plausible models of selection: one where low substitution rates or synonymous amino acid substitutions are favored (*i.e.*, purifying selection) and rates of nonsynonymous amino acids are not considered, and a second model calculating rates of both synonymous and nonsynonymous substitutions (*i.e.*, diversifying selection). A low (≤ 4.5) Bayes factor at a site indicates agreement with models favoring PPS, and a high (≥ 4.5) Bayes factor at a site indicates agreement with models allowing PDS. The second method used to detect evolutionary selection was a mixed-effects maximum likelihood model to detect episodic diversifying selection (EDS) at each amino acid site, and assumes that different strains are under variable selective pressures. Calculations were performed using the Mixed Effects Model of Evolution (MEME) model (41) on the DataMonkey 2.0 server (40). The likelihood ratio test (LRT) value for each site represents the level of agreement between two plausible models of selection: one allowing for the detection of nonsynonymous substitutions/diversifying selection, and a null model that does not. A higher (≥ 3) LRT value at a site indicates agreement with the model allowing EDS, and a lower (≤ 3) LRT value indicates agreement with the null model and suggests a lack of EDS at that amino acid position. We describe the calculated value at each site as the PDS, PPS, or EDS selection value (*i.e.*, the probability that the site is positively selected as reflected by deviation between full models and null models).

*Evolvability Calculations.* Evolvability values for ORF1a, S, and M proteins of the SARS-CoVs were calculated as previously described (31). Briefly, pairwise alignments using consensus amino acid sequences for ORF1a, S, and M of SARS-CoV and SARS-CoV-2 were generated using Clustal Omega (42). Pervasive selection values from Bayesian analyses were recoded for each aligned position. Positions falling within gaps between aligned segments were assigned a selection value of 1.0. Selection values for each amino acid position from the recently emerged SARS-CoV-2 were used as the base (expected values), and those from SARS-CoV were used as the query (observed values). Evolvability measures were not determined for the E, ORF3, ORF7, or ORF8 proteins because all sites were under purifying selection in both viruses, or in the ORF1b polyprotein or the N protein because multiple sites have already been subjected to pervasive diversifying selection in SARS-CoV and SARS-CoV-2.

*Selection of Traits for Analysis.* Nucleotide sequences encoding ORF1a, ORF1b, S, E, M, and N from SARS-CoV, SARS-CoV-2, and MERS-CoV were initially targeted for analysis. Sequences encoding ORF3, ORF7, and ORF8 from SARS-CoV and SARS-CoV-2 were also targeted. MERS-CoV genes sharing low homology to those in the SARS-CoV and SARS-CoV-2 genomes were not included in any analysis. Evolutionary selection analysis could not be performed on traits with insufficient diversity (< 3 distinct sequences) across the panel of strains (*i.e.*, E, ORF7). Evolvability analysis could not be performed on traits that are either strongly conserved or highly diverse across both the base and the query species (*i.e.*, ORF1b, N, E, ORF7, ORF8), and was therefore restricted to ORF1a, S, and M.

*Prediction of Functional Domains and N-Linked Glycans in Positively Selected or Evolvable Traits*. Functional domains were identified and localized for the ORF1a polyprotein, the ORF1b polyprotein, the S protein, the M protein, and the N protein from SARS-CoV, SARS-CoV-2, and MERS-CoV using the NCBI Conserved Domain Database (43). Prediction of sites amenable to N-linked glycosylation were made with GlycoEP using the binary profile of patterns algorithm (44).

*Protein Structure Predictions in Positively Selected or Evolvable Traits.* Three dimensional structure predictions were made for the ORF1a polyprotein, the ORF1b polyprotein, the S protein, the M protein, and the N protein from each species using PHYRE2 (45). High-confidence predictions were modeled as noted in Supplemental Table S2. In the absence of a high-confidence model for the M protein, a secondary structure prediction was made using GOR4 (46).

*Statistical Analysis.* Statistical tests were performed via the DataMonkey 2.0 server (40) or with GraphPad Prism 8.0.

*Data Availability.* All data relevant to this manuscript are presented in the main text and figures, are available in the Supplemental Materials, are posted at doi: 10.5281/zenodo.3756937, or can be accessed in public databases as noted in Supplemental Tables S1 – S3.
