## Supplemental Tables for "Selectomic and Evolvability Analyses of the Highly Pathogenic Betacoronaviruses SARS-CoV-2, SARS-CoV, and MERS-CoV"

**Supplemental Table S1: SARS-CoV2 Strain Information**

| **Strain** | **Accession** | **Location** | **Date** | **Source** | **DataBase** |
| --- | --- | --- | --- | --- | --- |
| Wuhan-Hu-1 | NC_045512 | China (Wuhan) | Dec-19 | sputum | GenBank |
| WHU02 | MN988669 | China (Wuhan) | 12/30/2020 | BAL fluid | GenBank |
| WHU01 | MN988668 | China (Wuhan) | Jan-02-2020 | BAL fluid | GenBank |
| USA-CA2/2020 | MN994468 | USA (CA) | Jan-22-2020 | NP swab | GenBank |
| HKU-SZ-005b_2020 | MN975262 | China | Jan-20 | sputum | GenBank |
| USA-WA1/2020 | MN985325 | USA (WA) | Jan-19-2020 | OP Swab | GenBank |
| USA-AZ1/2020 | MN997409 | USA (AZ) | jan-22-2020 | buccal swab | GenBank |
| USA-IL1/2020 | MN988713 | USA (IL) | Jan-21-2020 | sputum | GenBank |
| USA-CA1/2020 | MN994467 | USA (CA) | Jan-23-2020 | NP swab | GenBank |
| HKU-SZ-002a_2020 | MN938384 | China (Shenzen) | Jan-2020 | NP swab | GenBank |
| KCDC03 | EPI_ISL_407193 | South Korea | 1/25/2020 | OP Swab | GISAID |
| AI/I-004 | EPI_ISL_407084 | Japan (Aichi) | 1/25/2020 | OP Swab | GISAID |
| England02 | EPI_ISL_407073 | UK/England | 1/29/2020 | Swab (unsp) | GISAID |
| England01 | EPI_ISL_407071 | UK/England | 1/29/2020 | Swab (unsp) | GISAID |
| Singapore1 | EPI_ISL_406973 | Singapore | 1/23/2020 | sputum | GISAID |
| HZ-1 | EPI_ISL_406970 | China (Hangzhou) | 1/20/2020 | sputum | GISAID |
| BavPat1 | EPI_ISL_406862 | Germany (Bavaria) | Jan-20 | sputum | GISAID |
| VIC01 | EPI_ISL_406844 | Australia (Victoria) | 1/25/2020 | Unsp | GISAID |
| WH04 | EPI_ISL_406801 | China (Wuhan) | 1/5/2020 | BAL fluid | GISAID |
| WH03 | EPI_ISL_406800 | China (Wuhan) | 1/1/2020 | BAL fluid | GISAID |
| IDF0373 | EPI_ISL_40659 | France (Paris) | 1/23/2020 | OP Swab | GISAID |
| IDF0372 | EPI_ISL_406596 | France (Paris) | 1/23/2020 | OP Swab | GISAID |
| SZTH-004 | EPI_ISL_406595 | China (Shenzen) | 1/16/2020 | BAL fluid | GISAID |
| SZTH-003 | EPI_ISL_406594 | China (Shenzen) | 1/16/2020 | BAL fluid | GISAID |
| SZTH-002 | EPI_ISL_406593 | China (Shenzen) | 1/13/2020 | BAL fluid | GISAID |
| SZTH-001 | EPI_ISL_406592 | China (Shenzen) | 1/13/2020 | BAL fluid | GISAID |
| 20SF201 | EPI_ISL_406538 | China (Guangdong) | 1/23/2020 | Unsp | GISAID |
| 20SF211 | EPI_ISL_406536 | China (Foshan) | 1/22/2020 | Unsp | GISAID |
| 20SF210 | EPI_ISL_406535 | China (Foshan) | 1/22/2020 | Unsp | GISAID |
| 20SF207 | EPI_ISL_406534 | China (Foshan) | 1/22/2020 | Unsp | GISAID |
| 20SF206 | EPI_ISL_406533 | China (Guangzhou) | 1/22/2020 | Unsp | GISAID |
| 20SF174 | EPI_ISL_406531 | China (Zhuhai) | 1/22/2020 | Unsp | GISAID |
| Taiwan2 | EPI_ISL_406031 | Taiwan (Kaohsiung) | 1/23/2020 | Unsp | GISAID |
| HKU-SZ-002 | EPI_ISL_406030 | China (Shenzen) | 1/16/2020 | NP swab | GISAID |
| 20SF040 | EPI_ISL_403937 | China (Zhuhai) | 1/18/2020 | NP swab | GISAID |
| 20SF028 | EPI_ISL_403936 | China (Zhuhai) | 1/17/2020 | NP swab | GISAID |
| 20SF025 | EPI_ISL_403935 | China (Shenzen) | 1/15/2020 | OP swab | GISAID |
| 20SF014 | EPI_ISL_403934 | China (Shenzen) | 1/15/2020 | BAL fluid | GISAID |
| 20SF013 | EPI_ISL_403933 | China (Shenzen) | 1/15/2020 | ET aspirate | GISAID |
| 20SF012 | EPI_ISL_403932 | China (Shenzen) | 1/14/2020 | ET aspirate | GISAID |
| WZ-02 | EPI_ISL_404228 | China (Zhejiang) | 1/17/2020 | sputum | GISAID |
| WZ-01 | EPI_ISL_404227 | China (Zhejiang) | 1/16/2020 | sputum | GISAID |
| IPBCAMS-WH-03 | EPI_ISL_403930 | China (Wuhan) | 12/30/2020 | BAL fluid | GISAID |
| IPBCAMS-WH-04 | EPI_ISL_403929 | China (Wuhan) | 12/30/2020 | BAL fluid | GISAID |
| IPBCAMS-WH-05 | EPI_ISL_403928 | China (Wuhan) | 1/1/2020 | BAL fluid | GISAID |
| HBCDC-HB-01 | EPI_ISL_402132 | China (Wuhan) | 12/30/2020 | BAL fluid | GISAID |
| WIV07 | EPI_ISL_402130 | China (Wuhan) | 12/30/2020 | BAL fluid | GISAID |
| WIV06 | EPI_ISL_402129 | China (Wuhan) | 12/30/2020 | BAL fluid | GISAID |
| WIV05 | EPI_ISL_402128 | China (Wuhan) | 12/30/2020 | BAL fluid | GISAID |
| WIV02 | EPI_ISL_402127 | China (Wuhan) | 12/30/2020 | BAL fluid | GISAID |
| Nonthaburi74 | EPI_ISL_403963 | Thailand | 1/13/2020 | NP swab | GISAID |
| Nonthaburi61 | EPI_ISL_403962 | Thailand | 1/8/2020 | NP swab | GISAID |
| IVDC-HB-04 | EPI_ISL_402120 | China (Wuhan) | 1/1/2020 | BAL fluid | GISAID |
| IVDC-HB-05 | EPI_ISL_402121 | China (Wuhan) | 12/30/2020 | BAL fluid | GISAID |
| IVDC-HB-01 | EPI_ISL_402119 | China (Wuhan) | 12/30/2020 | BAL fluid | GISAID |

**Supplemental Table S2: SARS-CoV Strain Information**

| **Isolate** | **Accession #** | **Country** | **Isolation Date** | **Source** | **Ref PMID** |
| --- | --- | --- | --- | --- | --- |
| B039 | AY686864 | China | 2004 | Palm Civet |  |
| civet007 | AY572034 | China | 2004 | Civet | [16485471](https://www.ncbi.nlm.nih.gov/pubmed/16485471) |
| civet020 | AY572038 | China | 2004 | Civet | [16485471](https://www.ncbi.nlm.nih.gov/pubmed/16485471) |
| CUHK-W1 | AY278554 | Hong Kong | 2003 | Human | [12853594](https://www.ncbi.nlm.nih.gov/pubmed/12853594) |
| CV7 | DQ898174 | Canada | 2003 | Human | [14527350](https://www.ncbi.nlm.nih.gov/pubmed/14527350) |
| HKU-39849 | GU553363 | Hong Kong | 7-Jun-03 | Human |  |
| HSR 1 | AY323977 | Italy | 2003 | Human | [15109406](https://www.ncbi.nlm.nih.gov/pubmed/15109406) |
| Taiwan TC2 | AY338175 | Taiwan | 2003 | Human | [15812185](https://www.ncbi.nlm.nih.gov/pubmed/15812185) |
| Taiwan TC3 | AY348314 | Taiwan | 2003 | Human | [15812185](https://www.ncbi.nlm.nih.gov/pubmed/15812185) |
| TJF | AY654624 | China | 2004 | Swine | [15757562](https://www.ncbi.nlm.nih.gov/pubmed/15757562) |
| Tor2 | NC_028893 | Canada | 2003 | Human | 12730501 |
| TWY | AP006561 | Taiwan | 2003 | Human |  |
| Urbani | MK062179 |  | N/A | Human | [30393776](https://www.ncbi.nlm.nih.gov/pubmed/30393776) |
| ZJ0301 | DQ182595 | China | 2004 | Human | [14527350](https://www.ncbi.nlm.nih.gov/pubmed/14527350) |

**Supplemental Table S2: MERS-CoV1 Strain Information**

| **Isolate** | **Accession #** | **Country** | **Isolation Date** | **Source** | **REF PMID** |
| --- | --- | --- | --- | --- | --- |
| 2363 | MH395139 | Saudi Arabia | 4-Dec-16 | Human |  |
| 2366 | MH432120 | Saudi Arabia | 8-Feb-17 | Human |  |
| 011/DAB/C8/F<1 | MK357908 | Kenya | 21-Apr-17 | Camel N swab | [30482895](https://www.ncbi.nlm.nih.gov/pubmed/30482895) |
| 011/LOM/C20/F<1 | MK357909 | Kenya | 21-Apr-17 | Camel N swab | [30482895](https://www.ncbi.nlm.nih.gov/pubmed/30482895) |
| Amibara118/2017 | MK564474 | Ethiopia | 24-Aug-17 | Camel N swab | 31275264 |
| England2/2013 | KM015348 | UK | 10-Feb-13 | Human OP Swab |  |
| England4/2013 | KM210277 | UK | 13-Feb-13 | Human sputum |  |
| USA-2_Saudi Arabia_2014 | KP223131 | USA | June-1-2014 | Human sputum | 24827411 |
| FRA/UAE | KF745068 | France | 7-May-13 | Human sputum |  |
| Amman-Jordan-12918/2015 | MF000459 | Jordan | 7-Sep-15 | Human sputum |  |
| Artawiyah-KSA-13328/2016 | KX154694 | Saudi Arabia | 29-Feb-16 | Human sputum |  |
| Aseer-KSA-Rs924/2015 | KY688119 | Saudi Arabia | 1-May-15 | Human sputum |  |
| Hufuf-KSA-11002/2015 | KY688120 | Saudi Arabia | 10-May-15 | Human sputum |  |
| Jeddah-KSA-173RS1101/2017 | MN723542 | Saudi Arabia Jeddah | 1-Aug-17 | Human NP swab |  |
| Jordan-201440011123/2014 | MK039552 | Jordan | 21-Apr-14 | Human sputum |  |
| Khobar-KSA-6736/2015 | KY688118 | Saudi Arabia | 7-Feb-15 | Human sputum |  |
| Riyadh-KSA-18013832/2018 | MN723544 | Saudi Arabia | 30-Aug-18 | Human NP swab |  |
| FRA1_1627-2013_BAL_Sanger | KJ361500 | France | 26-Apr-13 | Human BAL fluid |  |
| KNIH/002_05_2015 | MK796425 | South Korea | 20-May-15 | Human |  |
| llama-passaged-Qatar1 | MN507638 | Qatar | N/A | Llama |  |
| Burkina Faso/CIRAD-HKU434 | MG923470 | Burkina Faso | 23-Feb-15 | Camel N swab | [29507189](https://www.ncbi.nlm.nih.gov/pubmed/29507189) |
| Ethiopia/AAU-EPHI-HKU4412 | MG923466 | Ethiopia | 15-Mar-17 | Camel N swab | [29507189](https://www.ncbi.nlm.nih.gov/pubmed/29507189) |
| Kenya/C1272/2018 | MH734115 | Kenya | 13-Mar-18 | Camel N Swab | [30570714](https://www.ncbi.nlm.nih.gov/pubmed/30570714) |
| Morocco/CIRAD-HKU213/2015 | MG923469 | Morocco | 3-Mar-15 | Camel N swab | [29507189](https://www.ncbi.nlm.nih.gov/pubmed/29507189) |
| Nigeria/NS004/2015 | MG923472 | Nigeria | 13-Jan-15 | Camel N swab | [29507189](https://www.ncbi.nlm.nih.gov/pubmed/29507189) |
| AHRI-FAO-1/2018 | MK967708 | Egypt | 2018 | Camel |  |
| KOR/KCDC/001_2018-TSVi | MK129253 | South Korea | 20-Oct-18 | OP Swab |  |
| KOR/KNIH/001_05_2015 | KT326819 | South Korea | May 22 2015 | Human sputum | 26473095 |
| Qatar15 | MK280984 | Qatar | 21-May-15 | Human | [31022948](https://www.ncbi.nlm.nih.gov/pubmed/31022948) |
| Riyadh_2_2012 | KF600652 | Saudi Arabia | 30-Oct-12 | Human NP Swab | [24055451](https://www.ncbi.nlm.nih.gov/pubmed/24055451) |

**Supplemental Table S4: Structural Modeling PHYRE Template Identifiers**

| **Query** | **Template** | **PHYRE Identifier** |
| --- | --- | --- |
| SARS-CoV-2 ORF1a | SARS-CoV ORF1a papain-like protease | c2fe8B |
| SARS-CoV-2 ORF1a | SARS-CoV ORF1a serine protease | d2duca1 |
| SARS-CoV ORF1a | SARS-CoV ORF1a papain-like protease | c2fe8B |
| SARS-CoV ORF1a | SARS-CoV ORF1a serine protease | d2duca1 |
| MERS-CoV ORF1a | MERS-CoV ORF1a papain-like protease | c4p16A |
| MERS-CoV ORF1a | MERS-CoV ORF1a serine protease | c2ynbA |
| SARS-CoV-2 ORF1b | SARS-CoV ORF1b NSP12 | c6nusA |
| SARS-CoV-2 ORF1b | SARS-CoV ORF1b NSP14 | c5c8sD |
| SARS-CoV-2 ORF1b | SARS-CoV ORF1b NSP16 | c3r24A |
| SARS-CoV ORF1b | SARS-CoV ORF1b NSP12 | c6nusA |
| SARS-CoV ORF1b | SARS-CoV ORF1b NSP14 | c5c8sD |
| SARS-CoV ORF1b | SARS-CoV ORF1b NSP16 | c3r24A |
| MERS-CoV ORF1b | SARS-CoV ORF1b NSP12 | c6nusA |
| MERS-CoV ORF1b | SARS-CoV ORF1b NSP14 | c5c8sD |
| MERS-CoV ORF1b | MERS-CoV ORF1b NSP16 | c5ynpA |
| SARS-CoV-2 S | SARS-CoV Spike | c5x5bB |
| SARS-CoV S | SARS-CoV Spike | c5x5bB |
| MERS-CoV S | MERS-CoV Spike | c5x5fC |
| SARS-CoV-2 N | SARS-CoV Nucleocapsid | d1sska |
| SARS-CoV N | SARS-CoV Nucleocapsid | d1sska |
| MERS-CoV N | MERS-CoV-1 Nucleocapsid | c4ud1B |
